# Solving the first novel protein structure by 3D micro-crystal electron diffraction

**DOI:** 10.1101/600387

**Authors:** H. Xu, H. Lebrette, M.T.B. Clabbers, J. Zhao, J.J. Griese, X. Zou, M. Högbom

**Affiliations:** Department of Materials and Environmental Chemistry, Stockholm University, 106 91 Stockholm, Sweden; Department of Biochemistry and Biophysics, Stockholm University, 106 91 Stockholm, Sweden; Department of Cell and Molecular Biology, Uppsala University, 751 24 Uppsala, Sweden

## Abstract

Micro-crystal electron diffraction (MicroED) has recently shown potential for structural biology. It enables studying biomolecules from micron-sized 3D crystals that are too small to be studied by conventional X-ray crystallography. However, to the best of our knowledge, MicroED has only been applied to re-determine protein structures that had already been solved previously by X-ray diffraction. Here we present the first unknown protein structure – an R2lox enzyme – solved using MicroED. The structure was phased by molecular replacement using a search model of 35% sequence identity. The resulting electrostatic scattering potential map at 3.0 Å resolution was of sufficient quality to allow accurate model building and refinement. Our results demonstrate that MicroED has the potential to become a widely applicable tool for revealing novel insights into protein structure and function, opening up new opportunities for structural biologists.

## Main Text

Electrons, similar to X-rays and neutrons, are a powerful source for diffraction experiments (Henderson, 1995). Due to the strong interactions between electrons and matter, crystals that are considered as powder in X-ray crystallography can be treated as single crystals by micro-crystal electron diffraction (MicroED (Shi *et al.*, 2013)). This enables structure determination of molecules from micron-to nanometer-sized 3D crystals that are too small for conventional X-ray diffraction (Shi *et al.*, 2013; Nannenga, Shi, Leslie *et al.*, 2014; Yonekura *et al.*, 2015; Clabbers *et al.*, 2017; Xu *et al.*, 2018). Furthermore, MicroED can be applied to study biomolecules of low molecular weight that are beyond what can be resolved by single particle cryo-EM imaging (Henderson, 1995; Khoshouei *et al.*, 2017).

Over the past decades, 3D ED methods have been developed for structure determination of small inorganic compounds (Mugnaioli *et al.*, 2009; Wan *et al.*, 2013; Gemmi *et al.*, 2015) and organic molecules (Kolb *et al.*, 2010; van Genderen *et al.*, 2016; Palatinus *et al.*, 2017). At the early stages of 3D ED method development, tilting of the crystal was done manually, while diffraction patterns were collected on negative film. It could take years before sufficient data were obtained and processed in order to determine the crystal structure (Zou *et al.*, 2003). The computerization of transmission electron microscopes (TEM) and the development of CCD detectors allowed software to be developed that can semi-automatically collect 3D ED data in less than an hour (Wan *et al.*, 2013; Kolb *et al.*, 2010). Thanks to the recent advancement in CMOS and hybrid detector technology, it is now feasible to collect diffraction data in movie mode while continuously rotating the crystal (Nannenga, Shi, Leslie *et al.*, 2014; Gemmi *et al.*, 2015; Nederlof *et al.*, 2013). Benefiting from these technological advances, data collection and structure refinement can now be performed within hours (Gruene *et al.*, 2018; Jones *et al.*, 2018). Furthermore, peptide structures have been solved *ab-initio* using high resolution MicroED data (Rodriguez *et al.*, 2015; de la Cruz *et al.*, 2017). Since 2013, several research groups have shown that it is feasible to re-determine already known protein structures using MicroED (table S1) and it is only very recently that a new polymorph of hen egg-white lysozyme was unveiled by MicroED, but again phased using a previously determined structure of the identical protein (Lanza *et al.*, 2019).

The sample handling of MicroED is similar to that of cryo-EM, while the data collection and processing are similar to that used in X-ray crystallography, making the technique highly adaptable to existing cryo-EM and general TEM labs (Nannenga, Shi, Leslie *et al.*, 2014; de la Cruz *et al.*, 2017; Shi *et al.*, 2013; Yonekura *et al.*, 2015; Clabbers *et al.*, 2017; Xu *et al.*, 2018). Therefore, it is of great benefit to further develop MicroED methods to meet the needs of scientists in a wide community.

Here, we report the next step in the development of MicroED by solving a novel protein structure, *Sulfolobus acidocaldarius* R2-like ligand-binding oxidase, *Sa*R2lox. The R2lox metalloenzyme family was discovered a decade ago from its similarities with the ribonucleotide reductase R2 protein (Andersson & Hogbom, 2009). Its physiological function is to date unknown, but the two crystal structures solved (Andersson & Hogbom, 2009; Griese *et al.*, 2013) of proteins belonging to this family reveal a 4-helix bundle core accommodating a dinuclear metal cluster, characterized by electron paramagnetic resonance (EPR) as Mn(III)/Fe(III) (Shafaat *et al.*, 2014), interacting with a long-chain fatty acid.

We demonstrate that MicroED data can be collected from *Sa*R2lox 3D micro-crystals by the continuous rotation method (Nederlof *et al.*, 2013; Nannenga, Shi, Leslie *et al.*, 2014). Conventional X-ray crystallography software can be used directly for processing ED data (*XDS* (Kabsch, 2010)), determining the phases using a homologous protein model of 35% sequence identity (*Phaser* (McCoy *et al.*, 2007)), and refining the model structure (*phenix.refine* (Afonine *et al.*, 2012)). These results illustrate that MicroED is a powerful tool for determining novel protein structures with sample requirements complementing those of X-ray crystallography, single particle cryo-EM and X-ray free electron lasers.

Micron-sized 3D crystals of *Sa*R2lox were grown using conventional hanging drop vapor diffusion, and are barely large enough to be distinguished under an optical microscope from precipitate and phase separation occurring in the drop (Fig. 1A). The mother liquor used for growing micro-crystals contains 44% polyethylene glycol 400. The viscosity of the sample prevented preparation of thin vitrified cryo-EM samples using the traditional deposit-blot-plunge routine. Thus, we used manual backside blotting to remove excess liquid and vitrified the sample in liquid ethane. The plate-like crystals are triangular in shape and a few micrometers in size (Fig. 1B). The thickness of the crystals is estimated to be less than 0.5 μm. We note that it is crucial to reduce the thickness of the protective layer of vitrified ice as much as possible in order to collect electron diffraction data of high signal-to-noise ratio from the crystals. The crystals diffracted beyond 3.0 Å in resolution as shown in Fig. 1C.

**Fig. 1.**
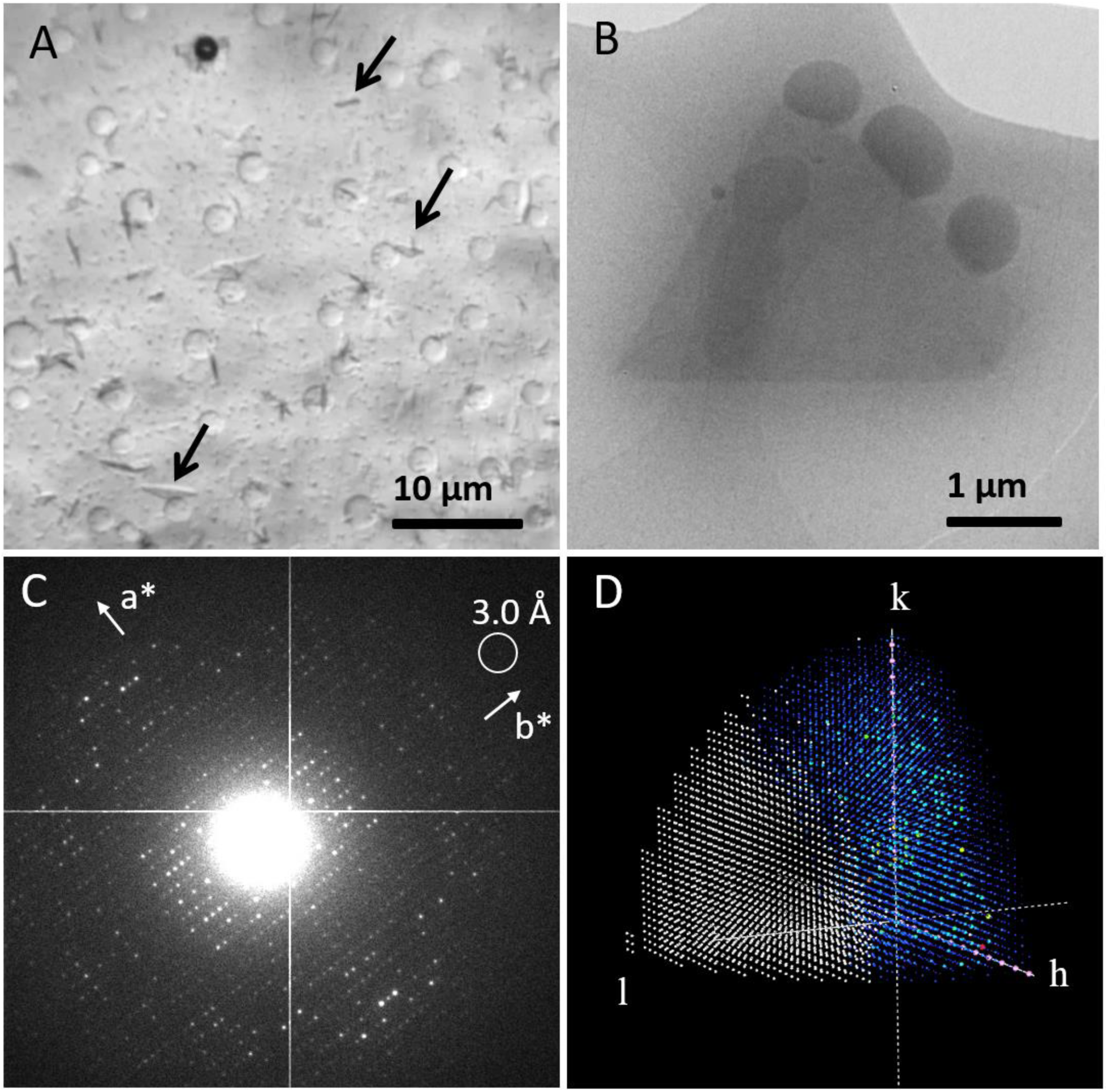
Overview of the MicroED experiment. (A) *Sa*R2lox micro-crystals (pointed out by arrows) viewed under optical microscope. (B) Over-focused TEM image of a typical diffracting *Sa*R2lox crystal preserved in vitrified ice. The volume of this particular crystal is estimated to be approximately 2 µm^3^, which contains approximately 6 × 10^6^ unit cells. (C) Typical electron diffraction pattern of the *Sa*R2lox crystal. The frame was taken after an accumulated electron dose of 4.3 e^-/Å2^. (D) Reconstructed reciprocal lattice showing limited data completeness with predominately missing reflections in the direction of *c*\*, missing observations are shown in white, observed reflections in rainbow color scheme, systematic absences are shown in pink.

MicroED data were collected by continuously rotating a single crystal in the electron beam at a rate of 0.45 °/second. The exposure time of each frame was 2 seconds, integrating over 0.90° of the reciprocal lattice. Data were typically collected over a total rotation range of 54° within 2 minutes. The electron dose rate applied was approximately 0.08 e^-^/Å^2^/s, resulting in a total dose of below 10 e^-^/Å^2^ per dataset. Because of their plate-like shape, a morphology commonly observed in protein crystallography, the *Sa*R2lox crystals are dispersed on the TEM grid with a preferred orientation. A total of 35 electron diffraction datasets were collected from the plate-like *Sa*R2lox crystals. Data were processed using crystallography software *XDS* (Kabsch, 2010). We could determine that the crystals are orthorhombic with a primitive unit cell using rotation electron diffraction processing software (REDp) (Wan *et al.*, 2013). Two screw axes were identified from the reflection conditions (Fig. 1C). Based on unit cell consistency and cross correlation, 21 out of the 35 datasets were merged. Due to the preferred orientation of the crystals, the data completeness increased from 50.9% for a single dataset to only 62.8% (Fig. 1D). However, the multiplicity (∼32) and overall I/σ(I) (6.12 up to 3.0 Å resolution) improved drastically (table S2). We demonstrated previously that the resulting structural model can be improved by merging data from a large number of crystals (Xu *et al.*, 2018).

The processed MicroED data were of sufficient quality to solve and refine the structure of *Sa*R2lox (Fig. 2A). The closest homologue to *Sa*R2lox in the Protein Data Bank (PDB) is an R2lox protein from *Mycobacterium tuberculosis* (*Mt*R2lox) sharing 35% sequence identity (PDB ID 3EE4 (Andersson & Hogbom, 2009)) (fig. S1). Using a modified search model based on the sequence alignment, we were able to phase the merged MicroED data by molecular replacement using *Phaser* (McCoy *et al.*, 2007), obtaining a clear single solution in space group *P*21212. The structure was iteratively built and refined in *COOT* (Emsley & Cowtan, 2004) and *phenix.refine* (Afonine *et al.*, 2012) (table S3). The refined electrostatic potential map provided sufficient detail to model side-chains and allowed rebuilding of the main chain (Fig. 2, B and D), even though certain parts of the maps were less well defined along the *c*\* direction (owing to the incomplete data). To assess the data quality and eliminate the influence of model bias, we generated a composite omit map covering the entire unit cell, confirming the correct interpretation for modelling the structure (Fig. 2, C and E).

**Fig. 2.**
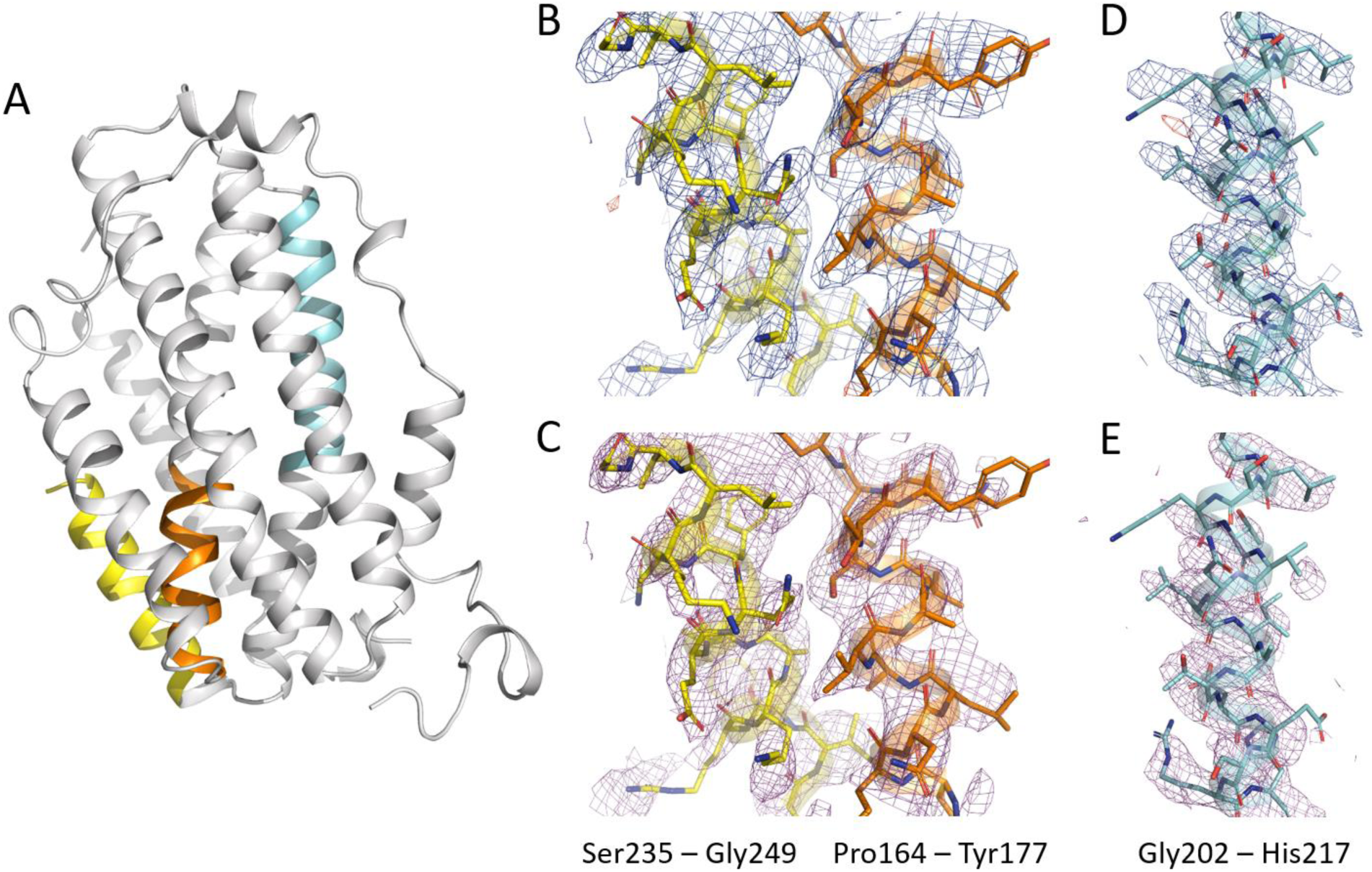
High quality electrostatic potential maps allowing accurate model interpretation. Overall structure of *Sa*R2lox solved by MicroED with three colored selections as examples for showing electrostatic potential maps in (B) to (E). Electrostatic scattering potential maps 2Fo-Fc (contoured at 1 σ, colored in blue) and Fo-Fc (contoured at ±3 σ, colored in green and red for positive and negative peaks, respectively) and simulated annealing composite omit maps (contoured at 1 σ, colored in magenta) are shown for residues 164 to 177 (orange) and 235 to 249 (yellow) in (B) and (C), respectively, and for residues 202 to 217 (cyan) in (D) and (E), respectively. Simulated annealing composite omit electrostatic potential maps are calculated with sequential 5% fractions of the structure omitted. Only observed reflections were used for map calculations, *i.e.* no missing F(obs) were restored using a weighted F(calc). Despite the low completeness, the data produces high quality well resolved maps. Oxygen and nitrogen atoms are colored red and blue, respectively, carbons are colored according to the selection previously mentioned.

The final model of *Sa*R2lox shows a protein backbone Cα root-mean-square deviation of 0.94 Å compared to the structure of *Mt*R2lox used for molecular replacement. This value is within expected range from proteins of this size and sequence identity level. The structure presented here confirms the dimeric biological assembly and the ferritin-like helix bundle overall fold, previously seen in the R2lox protein family (Andersson & Hogbom, 2009; Griese *et al.*, 2013) (fig. S2). Furthermore, the structure reveals new biochemically important features of the *Sa*R2lox enzyme, differing from the two structures known in this family: the substrate-binding pocket is reshaped and shows an altered electrostatic potential distribution (Fig. 3). In addition, the sequence identity with existing structures is lower than 40% which indicates a possibly divergent enzyme function (Tian & Skolnick, 2003). Thus, our results suggest that *Sa*R2lox has a different substrate specificity than the enzymes previously structurally characterized. Further biochemical studies are undergoing in our laboratory to confirm this hypothesis.

**Fig. 3.**
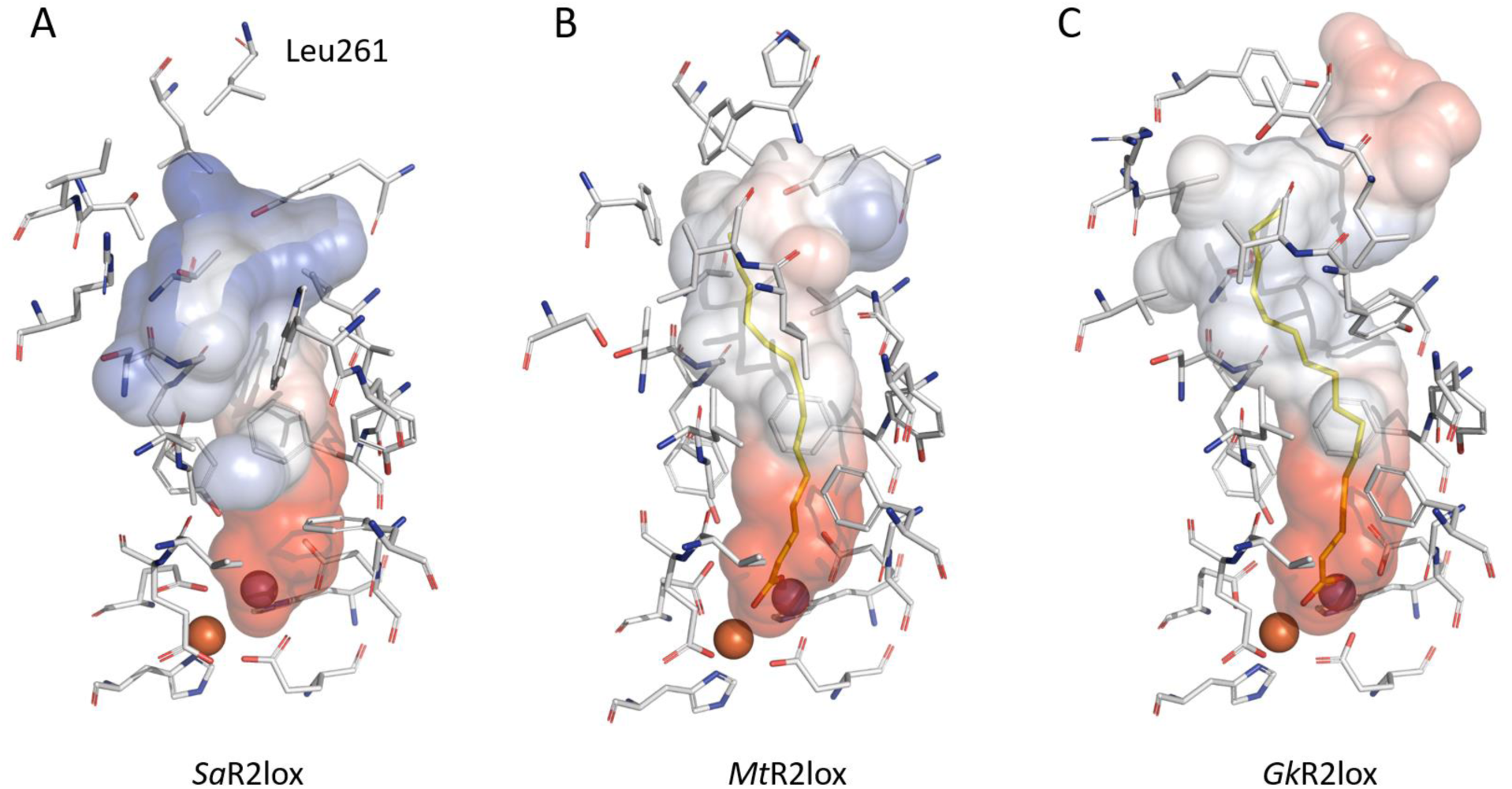
Differences between substrate-binding pockets support substrate specificity divergences in the R2lox metalloenzyme family. The molecular surface and identity of residues defining the substrate-binding pocket in the *Sa*R2lox structure (A) are compared with the two already known structures of this protein family, *Mt*R2lox (PDB ID 3EE4 (Andersson & Hogbom, 2009)) and (C) *Gk*R2lox (PDB ID 4HR0 (Griese *et al.*, 2013)). Although the Mn-and Fe-coordinating residues are fully conserved, the distal residues lining the cavity are remarkably divergent, impacting drastically the shape and electrostatic contact potentials of the pocket accommodating the putative substrate. This suggests that the substrate specificity could be different between these three enzymes, and thus their function could be divergent. Electrostatic contact potentials plotted on molecular surfaces are colored in a gradient from red (negative) to blue (positive). Ligands modelled as myristic acid and palmitic acid for *Mt*R2lox and *Gk*R2lox, respectively, are colored in yellow. A ligand was not modelled in the *Sa*R2lox structure due to a too weak signal in the corresponding scattering potential map but its presence cannot be excluded. Carbons, nitrogen, oxygen, manganese and iron atoms are colored in grey, blue, red, purple and orange, respectively. We note that the missing stretch of unmodeled residues in *Sa*R2lox (250 to 260) is not shared with *Mt*R2lox or *Gk*R2lox and is likely to impact distal closure of the cavity, further emphasizing differences between proteins.

MicroED data can be collected from crystals previously considered to be insufficient in size, which is its main advantage over X-ray crystallography. There are several aspects where electron diffraction can be further improved. For instance, by developing sample preparation methods such as cryogenic focused ion beam (Duyvesteyn *et al.*, 2018; Martynowycz *et al.*, 2019; Zhou *et al.*, 2019; Li *et al.*, 2018), the orientation of the crystals may be precisely controlled to achieve near 100% data completeness. Furthermore, the crystallographic quality indicators of the MicroED data are not yet as accurate nor as precise as generally achieved by X-ray crystallography. Rapid development of data collection strategies, as well as improvements of TEM hardware, may further accelerate the development of MicroED. Our results show that MicroED can be used for determining novel protein structures, even under the circumstances frequently encountered in macromolecular crystallography, where viscous sample environment, preferred orientation, and poor signal-to-noise ratio complicate data collection and structure determination.

## Acknowledgments

H.L. thanks P. Stenmark for discussions about this manuscript.

## Funding

We acknowledge financial support from the Knut and Alice Wallenberg Foundation through the project grants 3DEM-NATUR (no. 2012.0112, X.D.Z.) and Wallenberg Academy Fellows (no. 2017.0275, M.H.), the Science for Life Laboratory through the pilot project grant Electron Nanocrystallography, and the Swedish Research Council (2017-04018, M.H.; 2017-05333, H.Y.X.).;

## Author contributions

H.X contributed to project design, conception, electron diffraction data collection, electron diffraction data analysis, manuscript writing and figure making. H.L contributed to project design, crystal growth, structure determination, manuscript writing and figure making. M.C. contributed to electron diffraction data analysis, structure determination, manuscript writing and figure making. J.Z. contributed to electron diffraction data collection. J.G. contributed to cloning. X.Z. and M.H. contributed to project design, data analysis, conception and manuscript writing;

## Competing interests

The authors declare no competing interests;

## Data and materials availability

The atomic coordinates of the *Sa*R2lox structure are deposited in the Protein Data Bank under accession code 6QRZ. Raw MicroED data are available upon request. The python script used for converting diffraction frames from TIF to MRC and IMG (SMV format) is available upon request. REDp can be downloaded at mmk.su.se/zou.

## Supplementary Materials for

### Materials and Methods

#### Cloning, Expression and Protein Purification

A construct encoding full-length *Sulfolobus acidocaldarius* R2lox (accession number WP_011278976) was PCR-amplified from genomic DNA (DSM number 639, obtained from DSMZ) and inserted into pET-46 Ek/LIC (Novagen) using the following primers: GACGACGACAAGATGAAGGAAAAATTACTTGAATTCAGAAGT GAGGAGAAGCCCGGTTATTTGTCCAGCTTAATCTCCTCTATGAC.

Expression was carried out in *Escherichia coli* BL21(DE3) (Novagen). Cells were cultured at 37°C in terrific broth medium (Formedium) supplemented with 50 μg/ml ampicillin, in a bench-top bioreactor system (Harbinger). When an optical density at 600 nm of 0.7 was reached, expression was induced with 0.5 mM isopropyl β-D-1-thiogalactopyranoside (IPTG) and allowed to continue for 3 h. Then, cells were harvested by centrifugation and stored at -80 °C. The N-terminal His6 tagged *Sa*R2lox protein was purified via immobilized metal ion affinity chromatography and size-exclusion chromatography. Cells were resuspended in buffer A (25 mM Hepes-Na pH 7.0, 300 mM NaCl, and 20 mM imidazole) and disrupted by high-pressure homogenization. The lysate was cleared by centrifugation and applied to a nickel-nitrilotriacetic acid agarose (Protino) gravity flow column. The beads were washed extensively with buffer B (buffer A containing 40 mM imidazole). Protein was then eluted using buffer C (buffer A containing 250 mM imidazole), concentrated using Vivaspin 20 centrifugal concentrators with a 30,000 Da molecular weight cut-off polyethersulfone membrane (Sartorius), and applied to a HiLoad 16/60 Superdex 200 prep grade size-exclusion column (GE Healthcare) equilibrated in a final buffer of 25 mM Hepes-Na pH 7.0 and 50 mM NaCl. Fractions corresponding to the pure *Sa*R2lox protein were pooled, concentrated to 16 mg/mL, aliquoted, flash-frozen in liquid nitrogen and stored at -80 °C. The protein concentration was obtained using a calculated molecular weight for this construct of 38006 and an experimentally determined extinction coefficient at 280 nm for metal-bound protein of 52.13 mM^-1^.cm^-1^.

#### Crystallization

*Sa*R2lox protein was crystallized using the hanging drop vapor diffusion method. A volume of 2 µl of an 8 mg/ml protein solution is mixed with 2 µl of reservoir solution consisting of 44% (v/v) polyethylene glycol 400, 0.2 M lithium sulphate and 0.1 M sodium acetate (pH 3.4). Plate-like crystals grow within 48 hours at 21 °C.

#### Sample preparation

Cryo-EM sample of *Sa*R2lox is prepared by freezing the crystals in a thin layer of vitrified ice. A thin and uniform vitrified ice layer is crucial for obtaining MicroED of high signal-to-noise ratio. Meanwhile, the ice layer has to protect the crystals from being dehydrated under vacuum inside a transmission electron microscope (TEM). The 4 µl hanging drop is deposited onto a Quantifoil R 3.5/1 (300 mesh) Cu holy carbon TEM grid. The excessive liquid is removed by a manual back-side blotting. The grid is then rapidly plunge frozen in liquid ethane. We note that the automated blotting and vitrification routine using a FEI Vitrobot Mark IV was not efficient in removing the viscous mother liquid while leaving a sufficient number of crystals on the TEM grid.

#### Data collection

MicroED data were collected on a JEOL JEM2100 (LaB6 filament) TEM operated at 200 kV, with a Gatan 914 cryo-transfer holder. Before searching for suitable crystals, the electron beam is aligned, while the center of the TEM grid is brought to the mechanical eucentric height. During crystal searching, by inserting a 50 µm condenser lens aperture, the size of the electron beam is set to be slightly larger than the field of view (6 μm) on the side entry Orius detector. As soon as a suitable crystal is found, the beam is blanked while we set up for MicroED data collection. These measures are taken to avoid unnecessary electron dose on surrounding crystals. An area with a diameter of 2 µm as defined by a selected area aperture used to select the region of interest on the crystal. Electron diffraction data are collected by continuously rotating the SaR2lox crystal under the electron beam whilst simultaneously collecting the diffraction patterns on a fast Timepix hybrid pixel detector (Amsterdam Scientific Instruments). Data were collected at a sample to detector distance of 1830 mm, equivalent to 0.001198 Å^-1^ per pixel.

#### Data processing

The native format of collected data is TIF format. The data is converted to the Super Marty View (SMV) format using a python script developed in house. The important metadata of the experiment is written automatically to the headers of each SMV frame. Data were processed using XDS (Kabsch, 2010). The data were scaled and merged based on unit cell consistency, correlation coefficients between the datasets (analysed with *XDS nonisomorphism*) (Diederichs, 2017), I/σ(I) and resolution using *XSCALE* (Kabsch, 2010). Data were converted to MTZ format using *POINTLESS* (Evans & Murshudov, 2013), merged with *AIMLESS* (Evans, 2006), and structure factors were calculated using *TRUNCATE* (French & Wilson, 1978) Figure 1D was prepared with the Reflection Data Viewer in *Phenix* (Adams *et al.*, 2010).

#### Structure solution and refinement

The structure was solved by molecular replacement using *Phaser* (McCoy *et al.*, 2007) with atomic scattering factors for electrons. A truncated search model was created using the atomic coordinates of the R2-like ligand binding oxidase from *Mycobacterium tuberculosis* (PDB ID 3EE4, (Andersson & Hogbom, 2009)) using *Sculptor* (Adams *et al.*, 2010). A well-contrasted solution was obtained with one molecule per asymmetric unit in the space group *P*21212 (LLG = 161, TFZ = 14.0). The structure model was refined using rigid body refinement directly after molecular replacement in *phenix.refine* (Afonine *et al.*, 2012). The structure was iteratively built using *COOT* (Emsley & Cowtan, 2004), and refined using *phenix.refine* (Afonine *et al.*, 2012) with atomic scattering factors for electrons, automatic weighting of the geometry term and group B-factors per residue.

Certain regions of the map were less well resolved, in particular residues 250-260. As the map was difficult to interpret, we did not attempt to place those residues. Furthermore, a segment of an α-helix, residues 235-249, appeared to be shifted judging from the difference potential map after attempting to fit the corresponding side chains. This part was corrected and refined by real space refinement in *COOT* (Emsley & Cowtan, 2004) using geometrical restraints. Simulated annealing composite omit maps were generated using *phenix.composite_omit_map* (Afonine *et al.*, 2012), calculated with sequential 5% fractions of the structure omitted.

The model was validated using *MolProbity* (Chen *et al.*, 2010). Table S2 lists the crystallographic statistics in which the test set represents 5% of the reflections. The core root-mean-square deviation values between structures were calculated by the secondary-structure matching (SSM) tool (Krissinel & Henrick, 2004). Figures 2, 3, and S2 were prepared using the *PyMOL* Molecular Graphics System, version 2.2.3 Schrödinger, LLC. Electrostatic protein contact potentials were generated with the vacuum electrostatics tool in *PyMOL*, using the default values for cavity detection radius and cutoff, and using a level range of ± 50.

**Fig. S1.**
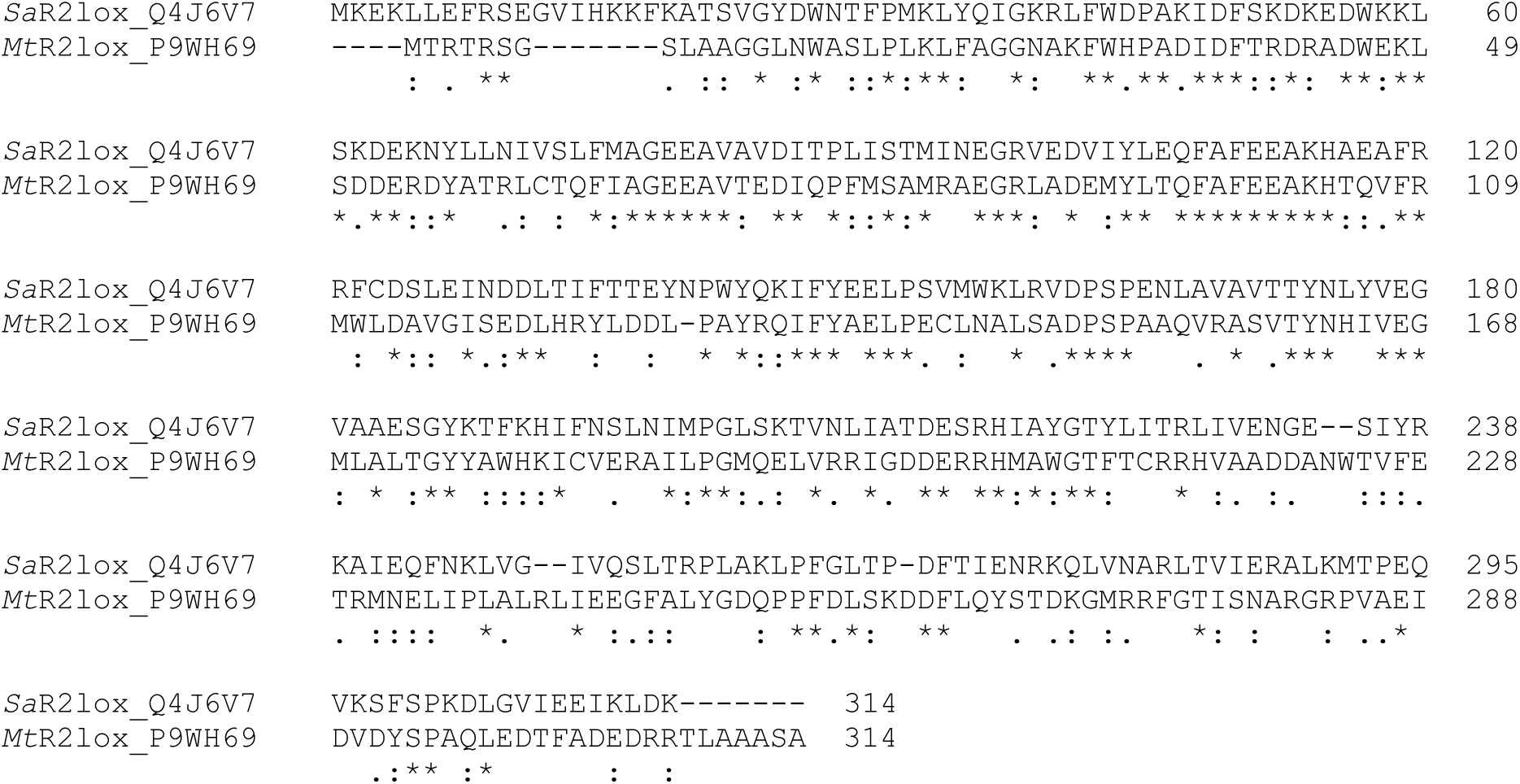
Sequence alignment between *Sa*R2lox (UniProt identifier Q4J6V7) and R2lox from *Mycobacterium tuberculosis* (*Mt*R2lox, UniProt identifier P9WH69) used as molecular replacement search model for the structure presented in this study (PDB ID 3EE4, (Andersson & Hogbom, 2009)) An asterisk symbol (*) indicates positions which have a single, fully conserved residue; a colon symbol (:) indicates conservation between groups of strongly similar properties; a period symbol (.) indicates conservation between groups of weakly similar properties. Sequence alignments were performed using ClustalW (2.1).

**Fig. S2.**
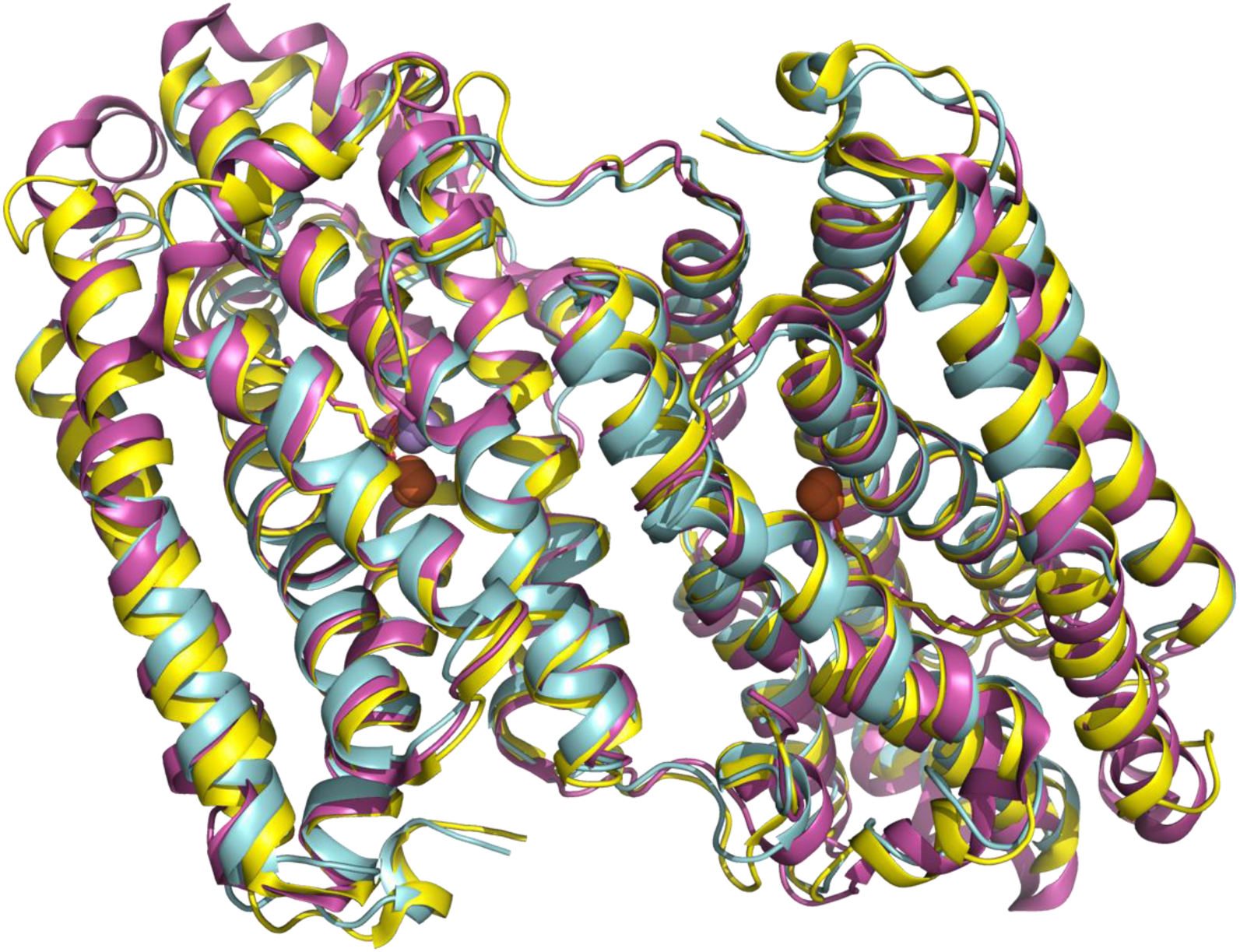
Overall superimposition of crystal structures of R2lox dimers. The two X-ray crystal structures of proteins from the R2lox family (PDB ID 3EE4 (Andersson & Hogbom, 2009) and 4HR0 (Griese *et al.*, 2013) colored in yellow and pink, respectively) are superimposed with the structure of *Sa*R2lox solved by MicroED in the current study (colored in cyan). The dimer assembly, generated by symmetry from the crystal packing, is the physiological oligomerization state of this family of proteins in solution. Manganese and iron ions are depicted as purple and orange spheres, respectively.

**Table S1.**
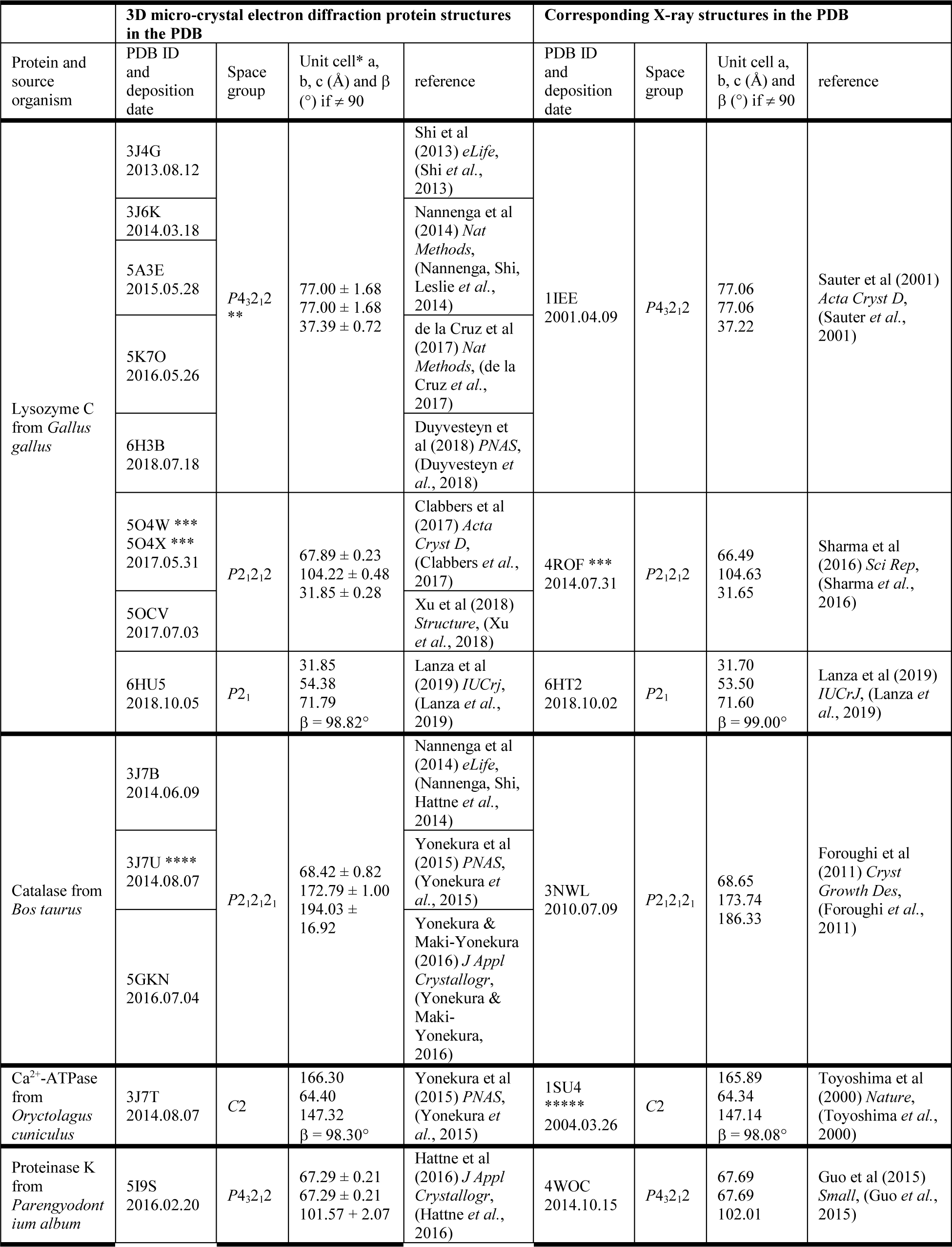
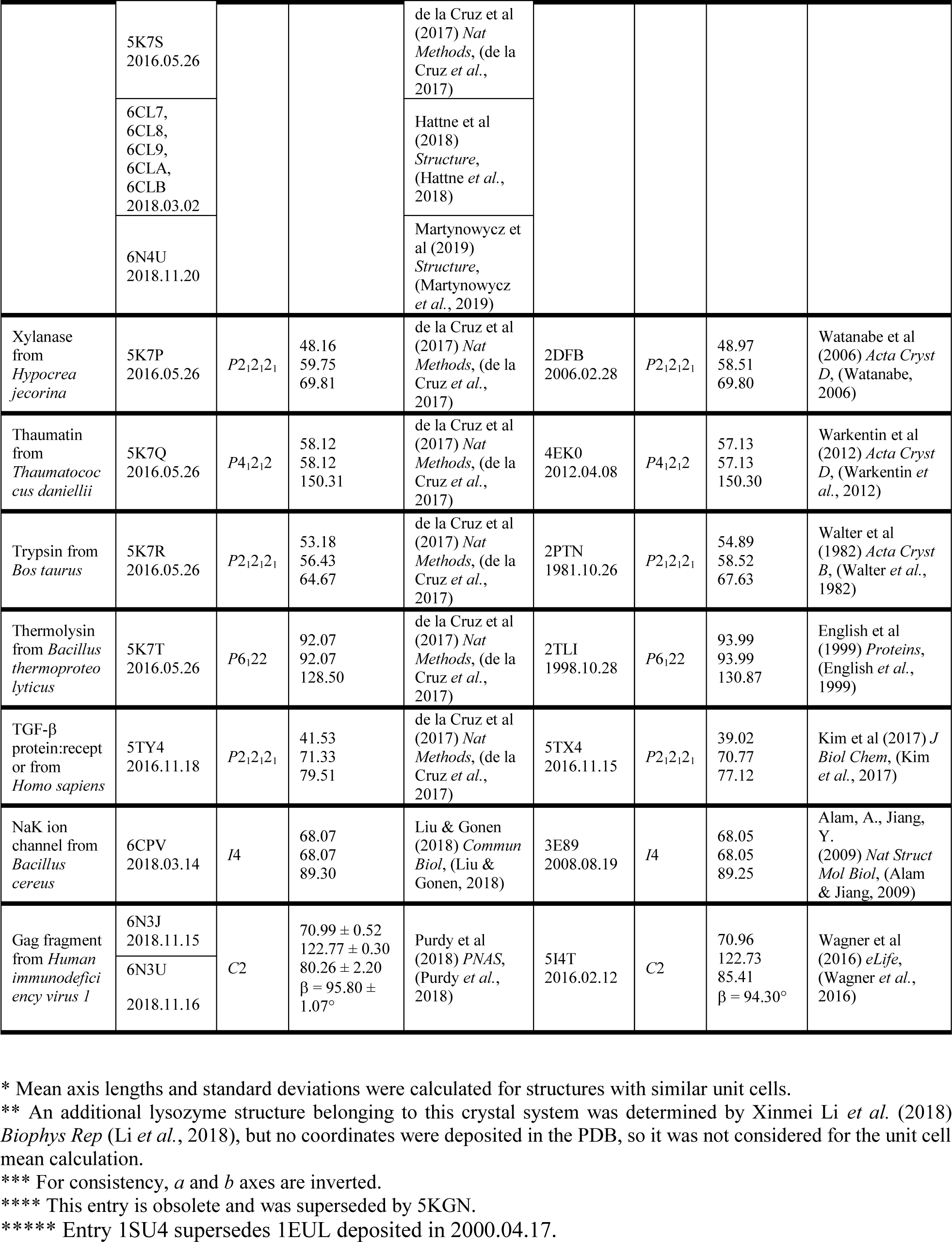
Overview of protein structures in the Protein Data Bank determined MicroED

**Table S2.**
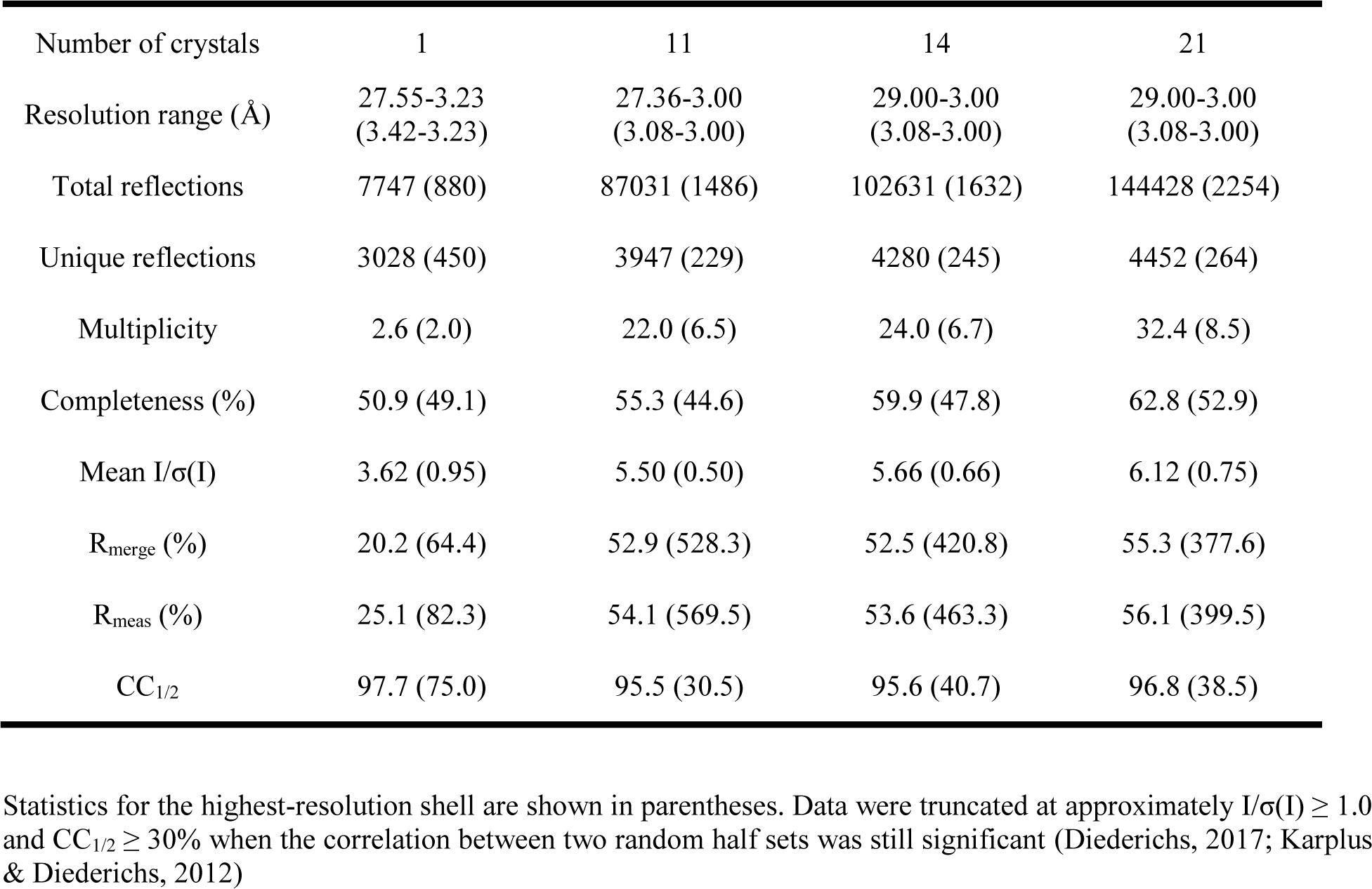
Electron diffraction merging statistics for a single crystal dataset, and merging of respectively 11, 14 and 21 crystal datasets.

| Number of crystals | 1 | 11 | 14 | 21 |
| --- | --- | --- | --- | --- |
| Resolution range (Å) | 27.55-3.23<br>(3.42-3.23) | 27.36-3.00<br>(3.08-3.00) | 29.00-3.00<br>(3.08-3.00) | 29.00-3.00<br>(3.08-3.00) |
| Total reflections | 7747 (880) | 87031 (1486) | 102631 (1632) | 144428 (2254) |
| Unique reflections | 3028 (450) | 3947 (229) | 4280 (245) | 4452 (264) |
| Multiplicity | 2.6 (2.0) | 22.0 (6.5) | 24.0 (6.7) | 32.4 (8.5) |
| Completeness (%) | 50.9 (49.1) | 55.3 (44.6) | 59.9 (47.8) | 62.8 (52.9) |
| Mean I/$\sigma$(I) | 3.62 (0.95) | 5.50 (0.50) | 5.66 (0.66) | 6.12 (0.75) |
| R<sub>merge</sub> (%) | 20.2 (64.4) | 52.9 (528.3) | 52.5 (420.8) | 55.3 (377.6) |
| R<sub>meas</sub> (%) | 25.1 (82.3) | 54.1 (569.5) | 53.6 (463.3) | 56.1 (399.5) |
| CC<sub>1/2</sub> | 97.7 (75.0) | 95.5 (30.5) | 95.6 (40.7) | 96.8 (38.5) |

Statistics for the highest-resolution shell are shown in parentheses. Data were truncated at approximately $I/\sigma(I) \geq 1.0$ and $CC_{1/2} \geq 30\%$ when the correlation between two random half sets was still significant (Diederichs, 2017; Karplus & Diederichs, 2012)

**Table S3.** Data collection and refinement statistics of *Sa*R2lox.

|  |  |
| --- | --- |
| **Data collection** |  |
| Wavelength (Å) | 0.02508 |
| Resolution range | 29.00-3.00 (3.08-3.00) |
| Space group | *P*2<sub>1</sub>2<sub>1</sub>2 |
| Unit cell dimensions |  |
| a, b, c (Å) | 63.31, 108.93, 48.17 |
| α, β, γ (°) | 90.00, 90.00, 90.00 |
| Total reflections | 144428 (2254) |
| Unique reflections | 4452 (264) |
| Multiplicity | 32.4 (8.5) |
| Completeness (%) | 62.8 (52.9) |
| Mean I/σ(I) | 6.12 (0.75) |
| Wilson B-factor | 47.93 |
| R<sub>merge</sub> | 0.553 (3.776) |
| R<sub>meas</sub> | 0.561 (3.995) |
| CC<sub>1/2</sub> | 0.981 (0.597) |
| CC* | 0.995 (0.860) |
| **Refinement** |  |
| Reflections used in refinement | 4423 (343) |
| Reflections used for R<sub>free</sub> * | 233 (15) |
| R<sub>work</sub> | 0.3179 (0.4299) |
| R<sub>free</sub> * | 0.3347 (0.4623) |
| Number of non-hydrogen atoms | 2243 |
| macromolecules | 2241 |
| ligands | 2 |
| solvent | - |
| Protein residues | 274 |
| RMSD(bonds) ** | 0.002 |
| RMSD(angles) ** | 0.41 |
| Ramachandran favoured (%) | 96.67 |
| Ramachandran allowed (%) | 3.33 |
| Ramachandran outliers (%) | 0.00 |
| Rotamer outliers (%) | 0.82 |
| Clashscore | 10.26 |
| Average B-factor | 48.37 |

Statistics for the highest-resolution shell are shown in parentheses. Data were truncated at approximately $I/\sigma(I) \geq 1.0$ and $CC_{1/2} \geq 50\%$ when the correlation between two random half sets was still significant (Diederichs, 2017; Karplus & Diederichs, 2012)
\* The R<sub>free</sub> test set represent approximately 5% of the reflections, the low number of reflections in the electron diffraction data test set might affect calculation of tabulated values
\*\* Root mean square deviation

